# Within and Across Species Variation in Plant Traits in Response to Long-Term Experimental Herbivore Exclusion in a Semi-Arid Shrubland

**DOI:** 10.1101/2024.06.24.600127

**Authors:** María del Pilar Fernández-Murillo, Fernando Daniel Alfaro, Douglas A. Kelt, Peter L. Meserve, Julio R. Gutiérrez, Alejandra J. Troncoso, Dylan Craven

## Abstract

1) Herbivores shape multiple facets of biodiversity patterns and ecosystem processes, yet are increasingly threatened by habitat loss and climate change. Here, we use a long-term experiment to examine how excluding herbivores, a proxy for biodiversity loss, restructures dominant plant ecological strategies within and across species.

2) We measured twelve functional traits of four dominant annual plant species in the semi-arid shrublands of Chile. We evaluated changes in trait distributions within and across species and changes in the coordination between above- and below-ground strategies in response to vertebrate herbivore exclusion.

3) Our study reveals three dominant ecological strategies characterizing trait variation within and across species, with an increase in the coordination between above- and below-ground strategies in the presence of herbivores. Our results show that herbivores induce shifts in ecological strategies, leading to conservative resource use, increased herbivore tolerance, reduced collaboration with soil microorganisms, and greater intraspecific diversity, facilitating species coexistence.

4) We empirically demonstrate how herbivores shape ecological strategies within and across species, and demonstrate that the absence of herbivores alters the diversity of adaptation strategies of plants and leads to a decoupling of resource acquisition strategies above- and below-ground. These results advance understanding of annual plant responses to herbivore loss.

## INTRODUCTION

The loss of biodiversity is a critical driver of global change, with the absence of specific groups and guilds, such as herbivores, exerting profound impacts on ecosystem functioning (Roy et al., 2020). Numerous studies have highlighted that plants undergo adaptive changes in response to herbivore pressure, leading to shifts in ecological strategies to ensure survival (Roy et al., 2020; Rauschkolb et al., 2022). These strategies are intricately linked to functional traits, which may vary due to the influence of environmental conditions or conspecific and interspecific interactions (Funk et al., 2016).

Due to their short lifespan, annual herbaceous plants are generally considered to adapt rapidly to disturbances (Grime 1979), making them an ideal model for examining changes in ecological strategies (Mulroy & Rundel 1977; Chesson et al., 2000; Funk et al., 2021; Rauschkolb et al., 2022). Especially in water-limited environments, they play a crucial role in food webs, contributing a significant portion of biomass and plant diversity (Li et al., 2008). Nevertheless, most studies addressing the relationship between herbivory and shifts in plant ecological strategies have largely focused on shrubs or perennial herbs (Archibald et al.,2019). Therefore, understanding the diversity of ecological strategies that annual plants employ in the absence of herbivores could be crucial for predicting changes in ecosystem functioning due to global change.

Above-ground trophic interactions drive changes in functional traits, from leaf nutrient content to root structure (Gundale & Kardol 2021). For instance, loss of foliage directly impacts the quality and quantity of plant tissues and resources (Bardgett & Wardle 2003), while the resulting decrease in productivity and increase in carbon allocation to secondary defenses may negatively impact plant growth (Peschiutta et al., 2018). On the other hand, herbivores could have compensatory effects on plant growth and coexistence through the reallocation of resources and nutrients to soils (Roy & Bagchi, 2022; Roy et al., 2022), and they may regulate dominant or generalist species (Mortensen et al., 2018). Variation in functional traits, primarily attributed to resource availability, can be strongly influenced by additional ecological interactions such as herbivory (Cipollini, 2004; Violle et al., 2009).

Key traits such as plant height and leaf area, essential for light competition and plant-herbivore tolerance strategies, also respond to gradients in (Zhang et al., 2021). Similarly, traits such as leaf nitrogen content and specific leaf area (SLA) can reflect resource availability and growth potential of plants, and their variation underlies plant-herbivore tolerance strategies (Laliberte et al., 2012).

Herbivores alter habitat conditions in diverse ways, affecting plant community composition and structure by altering resource availability and modifying competitive relationships (Chesson 2000). Under resource limiting conditions, competition may lead to greater niche differentiation; competing species can coexist only if they differ in trait values (“limiting similarity”; MacArthur & Levins 1967; Emery, 2007; Lohbeck et al., 2013; Kraft et al., 2015). In the absence of herbivory, lower SLA and higher maximum plant height may enable greater resource conservation as a response to resource limitation (Westoby et al., 2002; Reich, 2014). However, in the presence of herbivores, vegetation cover typically diminishes, leading to increased soil exposure. Consequently, there is higher light intensity at the soil surface, elevated drought stress, and an accumulation of soil nutrients, attributable to the greater input of organic material and high rates of decomposition.

Drought stress may increase photosynthetic rates (Kirschbaum, 2011), and therefore values of other traits also may be associated with a resource acquisition strategy (Reich 2014). The alteration of soil structure and nutrient availability caused by herbivory also affects below-ground features. For example, reduced nutrient availability in the soil, caused by lower plant biomass and greater soil compaction, will be reflected in the increased association of plants with bacteria to facilitate access to nutrients (Zhang et al., 2021). Conversely, increases in soil removal due to belowground herbivory may facilitate resource acquisition, which is reflected in increased root length and biomass (Roumet et al., 2016; Zhang et al., 2021). Other root traits associated with a resource acquisitive strategy include high specific root length (SRL), specific root surface area (SRSA), and low root diameter (RD; Fort et al., 2013; Ravenek et al., 2016).

The response of plants to herbivore pressure may be less evident at the community level because species may respond differently, highlighting the need to evaluate community-wide responses to herbivory (Kimball et al., 2016). Intraspecific variation also can drive community trait responses to environmental variation. For instance, intraspecific variation accounted for up to 44% of variation in the community-weighted mean (CWM) of several key functional traits along an elevation gradient in a floodplain grassland community (Jung et al. 2010). Similarly, Kichenin et al. (2013) found weak SLA responses, and decreasing leaf nitrogen (N) and phosphorus (P) concentrations, along an altitudinal gradient.

However, these patterns resulted from contrasting responses between species, which are frequently overlooked in species-based measures of plant community responses (Violle et al. 2012). On the other hand, some traits, such as leaf mass area (LMA; 1/SLA), may have greater intraspecific variation than others, such as wood density (WD; Fajardo & Piper 2011). While there is growing evidence of the importance of accounting for intraspecific trait variation in quantifying community-level responses to resource gradients (Siefert et al. 2015), the extent to which herbivores affect intraspecific trait variation is uncertain.

Across spatial scales, trait variation within and across plant species can be classified into non-mutually exclusive ecological strategies. The Leaf Economics Spectrum (LES) elucidates how traits associated with resource-use will influence growth rates and ecological filtering along resource gradients (Wright et al., 2007; Reich, 2014). Whereas acquisitive species should exhibit high growth rates under resource-rich conditions, conservative species are expected to minimize performance losses at the expense of lower growth rates. These divergent responses should, in turn, lead to ecological filtering along resource gradients (Reich, 2014). The Herbivory Tolerance Spectrum (HTS) integrates trait values that could indicate a higher tolerance to direct consumption of plant tissues by herbivores. For example, since leaves with higher carbon content and higher toughness are less palatable than leaves with higher energy (e.g., nitrogen) content, the former would be selected for under conditions of high herbivory pressure. Similarly, herbivory pressure should lead to shorter, more shrub-like plants. In contrast, in the absence of herbivory, trait values such as taller plants and leaves with more nitrogen content should be more frequent (Carmona et al., 2011). The Root Economics Spectrum (RES) parallels the Leaf Economics Spectrum (LES), reflecting how root traits are linked to below-ground resource acquisition, including traits like root length and specific root length (SRL) that are associated with resource availability (Prieto et al., 2015; Roumet et al., 2016). A complementary hypothesis to the RES is the Collaboration Spectrum (CS) (Bergman et al., 2020), which associates high specific root length (SRL) in short-lived species with a "do-it-yourself" strategy for water and nutrient acquisition, while a low SRL and large root diameters indicate a greater dependency on mycorrhizal fungi for efficient resource acquisition.

Above- and below-ground traits of plants are intrinsically interconnected and respond jointly to variation in environmental conditions (Craine & Towne 2010). Previous studies have highlighted the importance of coordination between roots and leaves for the acquisition of resources such as light and water (Freschet et al., 2015; Liu et al., 2010). This coordination among traits improves the ability of plants to acquire scarce resources, or reduces their dependence on specific resources (Freschet et al., 2018). Nevertheless, the existence of coordination between above- and below-ground traits remains unclear, given the considerable variation observed across different ecosystems and environmental conditions (e.g., Fort et al., 2013; Liu et al., 2010; Carvajal et al., 2019; Weigelt et al., 2021). Therefore, characterizing coordination among plant traits is crucial for deepening current understanding of the potential impacts of global change drivers on plant communities and ecosystem processes (Weigelt et al., 2021).

We evaluate the long-term impacts of herbivore loss on trait diversity within and across species of annual plants in a semi-arid ecosystem. We address the following questions: How does the long-term absence of herbivory affect ecological collaboration strategies, herbivory tolerance, and resource acquisition within and across annual plant species? Furthermore, to what extent does the loss of herbivores impact the coordination between above- and below-ground resource acquisition strategies? We expect that in the absence of herbivores, above- and below-ground traits will exhibit similar and complementary responses, characterized by more conservative, less collaborative, and less herbivore-tolerant phenotypes (Diaz et al., 2007; Rota et al., 2017; Blumenthal et al., 2020). We further expect that, in the absence of herbivores, trait diversity of above- and below-ground traits will be greater across species than within species (Siefert et al. 2015).

## METHODS

### Study area

We conducted this study at the Long-Term Socio-Ecological Research (LTSER) site in Bosque Fray Jorge National Park in north-central Chile (-30.655 Lat.; -71.682° Long.). Established in 1989, this experiment consists of twenty 75 x 75 m plots randomly assigned to five treatments, each with four replicates (Gutiérrez et al., 2010; Kelt et al., 2013). The present study focuses on the control and herbivore exclusion treatments, hereafter referred to as with and without herbivores, respectively. The without herbivores treatment was established in 2007 and consists of a 2 m tall steel mesh fence buried at a depth of 50 cm. These fences prevent the entry of small mammals, which are the most abundant vertebrate herbivores in the study site area, as well as non-native European rabbit and European hare (*Oryctolagus cuniculus* and *Lepus europeaus*, respectively). The with herbivores treatment does not have physical barriers, thereby allowing small mammals and non-native lagomorphs to move freely. Herein, we analyze data on the annual plant communities in these two treatments in 2021, following 14 years of herbivore exclusion.

The climate of the study site is Mediterranean, characterized by austral fall and winter precipitation pulses ranging from 0.5-50 mm (Ulbrich et al., 2012). In rainy years, mean annual precipitation is greater than 150 mm, while in dry years it does not exceed 50 mm (CEAZA, 2023). In 2021, the total annual precipitation was 19.7 mm, and the average annual temperature was 13.4 °C. (Figure S1; CEAZA, 2023). Total precipitation in 2021 was among the driest years on record due to the ongoing megadrought in central Chile (Garreaud et al., 2020), which may have affected the number and composition of species sampled for this study.

### Study species and communities

Since the beginning of the study, we have recorded a total of 51 annual plant species, most of which occur in wet years. In 2021, the year of this study (Fig. S1), we recorded 14 species in control plots, and 13 in herbivore exclusion plots. To evaluate the influence of herbivory on intraspecific trait variation, we selected plant species found in both treatments; only seven species were present in both treatments, of which three were geophytes. We do not consider geophyte species as these species may have ecological strategies that are not directly comparable with annual plants. The four annual plant species that occurred in both experimental treatments were: *Bromus berteroanus*, *Moscharia pinnatifida, Plantago hispidula* and *Viola pusilla* (Fig. S2 A-D).

### Functional traits

We collected individuals for trait sampling in August 2021, when annual species are typically most abundant (Madrigal et al., 2016; Fernández-Murillo et al., 2023). We measured above- and below-ground traits of 224 individuals, which consisted of 28 adult individuals per species and treatment (N = 28 individuals x 4 species x 2 treatments). For each individual, we measured 13 functional traits (Table S1), which we selected because they have been associated previously with an array of plant ecological strategies (e.g., Díaz et al. 2016, Bergmann et al. 2020): plant height (HT), leaf thickness (LT), specific leaf area (SLA; cm^2^ g ^−1^), and leaf dry matter content (LDMC; g g^-1^) following standardized protocols (Pérez-Harguindeguy et al., 2016). We used a vernier caliper to measure leaf thickness and plant height in the field and stored leaves in plastic bags with a wet paper towel in a cooler to minimize transpiration. Within approximately 24 hours, we weighed leaves to obtain fresh mass with a digital balance (±0.1mg) and scanned them with a digital scanner at a resolution of 1200 DPI (HP Smart Tank model 619). We oven-dried leaves at 70 °C for 72 hours and weighed them to determine dry mass. We measured leaf area (cm^2^) using ImageJ (Rasband 1997). We calculated SLA by dividing leaf area by dry leaf mass and LMDC by taking the difference between fresh and dry leaf mass divided by fresh mass. We also measured C and N concentrations for all leaf tissue samples and the isotopic composition of carbon (δ^13^C) and nitrogen (δ^15^N) using a Thermo Delta V Advantage IRMS, coupled with a Flash2000Elemental Analyzer (Waltham, Massachusetts, USA).

We extracted all individuals from the soil to measure below-ground traits. We carefully removed the soil to a depth of approximately 5 cm for each individual, and gently pulled them up from the base to avoid damaging fine roots, and cleaned their roots by hand. We oven-dried samples at 70°C for 72 hours, weighed them, and scanned them using the same procedure as for leaves. Using the program RhizoVision (Seethepalli et al., 2021), we measured root length, surface area, and average diameter, which we used to calculate the following below-ground traits: root diameter (RD; mm), specific root length (SRL; cm g^-1^), and specific root surface area (SRA; cm^2^ mg^-1^). We classified roots of the first and second orders based on their diameter, with roots having a diameter less than 0.2 mm being classified as fine roots (McCormack et al., 2015). We then calculated the percentage of fine roots (based on length) as a proxy for the absorptive capacity of the root system.

### Data analysis

Prior to analysis, we normalized trait data by logarithmically transforming variables to meet normality assumptions as necessary and applying z-score standardization to all variables. To identify above- and below-ground ecological strategies and to detect if they shifted across treatments, we performed a principal component analysis (PCA) using the “PCA” function in the R package “vegan” (Oksanen et al., 2017), the analysis was performed with a covariance matrix. We then fitted a PERMANOVA to test for statistical differences in trait spaces between treatments, species, and their two-way interaction with the “*adonis”* function in the R package ’vegan’ (Oksanen et al., 2017). We compared the distributions of the first two principal components obtained in the PCA for above- and below-ground traits separately and together for the annual plant communities (level interspecific) and each species (intraspecific) (Fig. 1; Capdevila et al., 2023). We assigned the following ecological strategies to components of both PCAs: 1) the *Leaf Economics Spectrum (LES)* captures the trade-off between resource acquisition and conservation (Wright et al. 2004, Reich 2014, Diaz et al., 2016), which was driven principally by variation in HT, SLA, LDMC, and LT; 2) the *Herbivory Tolerance Spectrum (HTS)*, which reflects variation in investment in mechanically and chemically protecting leaves and roots from herbivory; 3) the *Root Economy Spectrum (RES)*, which is similar to that described for above-ground traits (e.g., LES), but which captures the trade-off between resource acquisition and resource conservation for below-ground traits, and is principally associated with variation in RL and RDMC; and lastly 4) the *Collaboration Spectrum (CS)*, which describes variation in below-ground traits, principally SRL, that reflect how plants acquire water and nutrients, either directly, i.e. a “do-it-yourself” strategy, or in collaboration with mycorrhizal fungi (Fig.1, Table S1).

**Figure 1.**
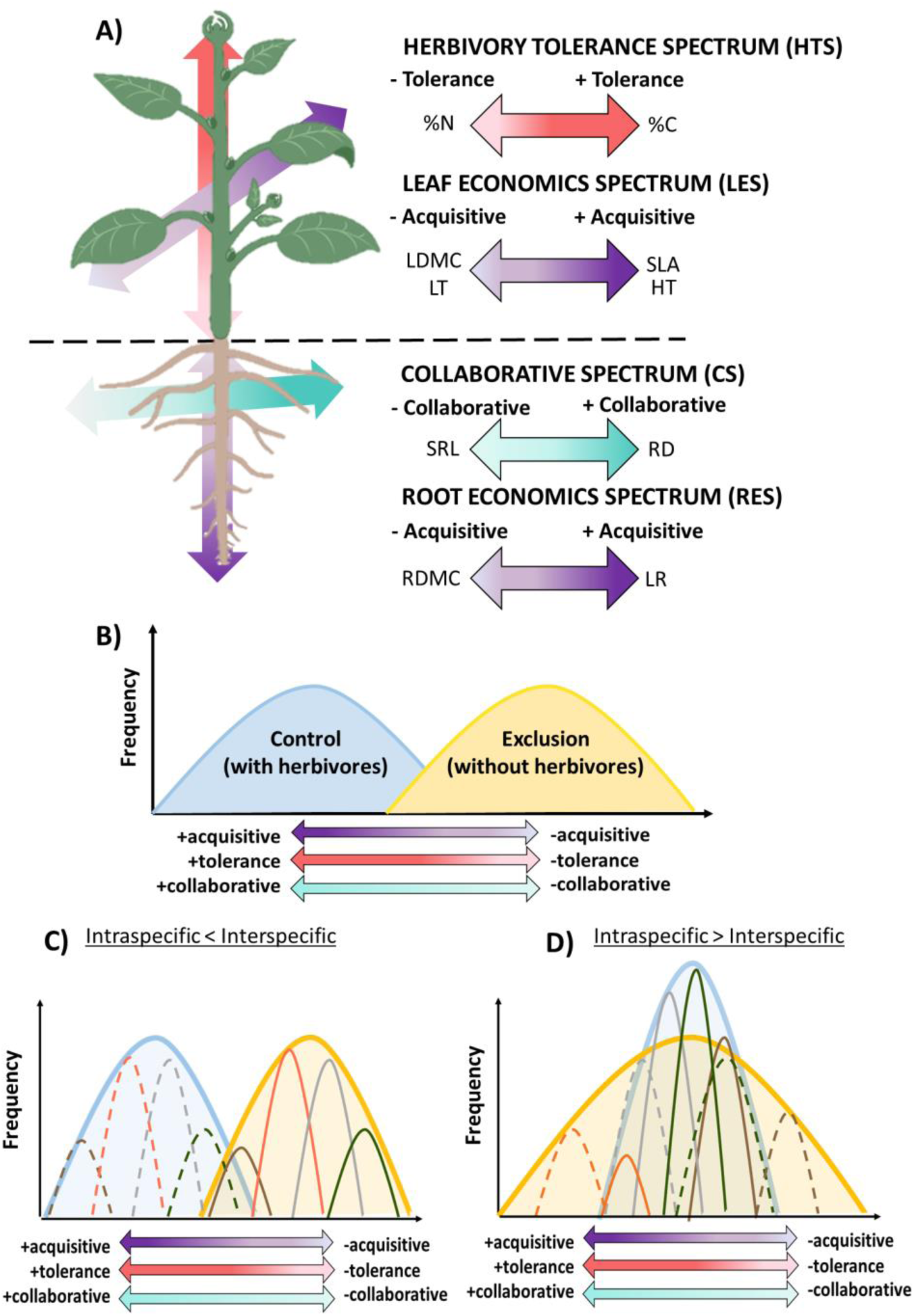
A) We integrate above- and below-ground functional traits into the conceptual frameworks of the Leaf Economics Spectrum (LES) and the Root Economics Spectrum (RES) with the Herbivory Tolerance Spectrum (HTS) and Collaboration Spectrum (CS) spectra. We anticipate that variation in above- and below-ground traits will be coordinated among annual plant species that may interact with herbivores (control treatment), yet will be decoupled among annual plant species that do not interact without herbivores (herbivore exclusion treatment). B) Additionally, we expect that plants in plots with herbivores will exhibit more acquisitive, collaborative, and herbivory tolerant trait values than plants in plots without herbivores. Lastly, the impacts of herbivory may scale up to the community level by either: C) lowering intraspecific trait variation, leading to greater interspecific trait variation, and differences in community-level trait means between plots with and without herbivores, or D) increasing intraspecific trait variation, which will decrease community-level trait differences between communities with and without herbivores. In C and D, each colored curve represents different species within the community. The segmented curves depict the same species as the solid curves; however, the former indicates the distribution of a ecological strategy in a herbivore-present community (blue), whereas the latter depicts the distribution of a ecological strategy in a community without herbivores (yellow).

For each spectrum (dimension), we calculated the diversity of ecological strategies using four statistical moments of their distributions, i.e., mean, variance, skewness, and kurtosis (Enquist et al., 2015; Maitner et al., 2023). The mean represents the dominant ecological strategy within communities or species (Violle et al., 2012; Maitner et al., 2023). Variance is a proxy for the breadth of niche (Gaedke & Klauschies, 2017; Maitner et al., 2023), as it measures how ecological strategies are distributed across species and communities.

Skewness refers to the degree to which the distributions of strategies ecological are balanced or unbalanced; higher absolute values indicate the presence of a dominant strategy or a higher abundance of individuals with exceptional trait values, which is often caused by asymmetric competition or rapid environmental changes (Enquist et al., 2015; Maitner et al., 2023). Kurtosis reflects the evenness of distributions of ecological strategies; high values reflect a narrow distribution and suggest that many species or individuals with similar ecological strategies co-occur, indicating low diversity of ecological strategies, possibly due to environmental filtering. In contrast, low kurtosis values reflect a broader distribution, indicating that ecological strategies are evenly distributed, indicating high diversity of ecological strategies, perhaps due to limiting similarity (Cornwell & Ackerly, 2009; Maitner et al., 2023).

To estimate the statistical moments of the distributions of PC1 and PC2 of the above- and below-ground ecological strategies at the community level (i.e., across all species and individuals; “interspecific”) and for each species (across all individuals in a species; “intra-specific”), we used a bootstrapping approach using the function “trait_np_bootstrap” and 1000 bootstrap replicates with the R package “traitstrap” (Maitner et al., 2023). We interpreted differences between treatments or species as significant if their 95% confidence intervals did not overlap. To test the coordination of resource economy strategies between above- and below-ground traits, we performed a Pearson correlation analysis between the PC2 dimensions of the PCA of traits of above-ground and below- ground components, both corresponding to resource economy strategies, respectively. All analyses and visualization were done using R version 4.3.0 (R Core Team 2023).

## RESULTS

### Above- and below-ground ecological strategies

The first two components of the PCA of above-ground traits jointly explained 49.8% of total variance (Fig. 2A; Tables S2 and S3). The first principal component explained 26.7% of the variance, capturing variation in the herbivory tolerance gradient as it is correlated with leaf C/N, %N, and %C in the leaf (Fig. 2A; Table S3). On the left extreme of this gradient, individuals with greater herbivory tolerance have high leaf %N, and low %C, while those at the other extreme (right) have high %C and low %N. The second principal component explained 23.1% of the variance and captures variation in resource acquisition, as it is correlated with HT, δ^15^N, and SLA (Fig. 2A; Table S3). At the lower extreme of the resource acquisition gradient, individuals with high SLA, HT, and low δ^15^N have values associated with resource acquisition, while those at the upper extreme have values associated with resource conservation, i.e., high LT and LDMC (Fig. 2A). PERMANOVA analysis indicates statistically significant differences in trait spaces between species (P < 0.0001) and experimental treatments (P < 0.01), but not for the interaction of both (P > 0.05).

**Figure 2.**
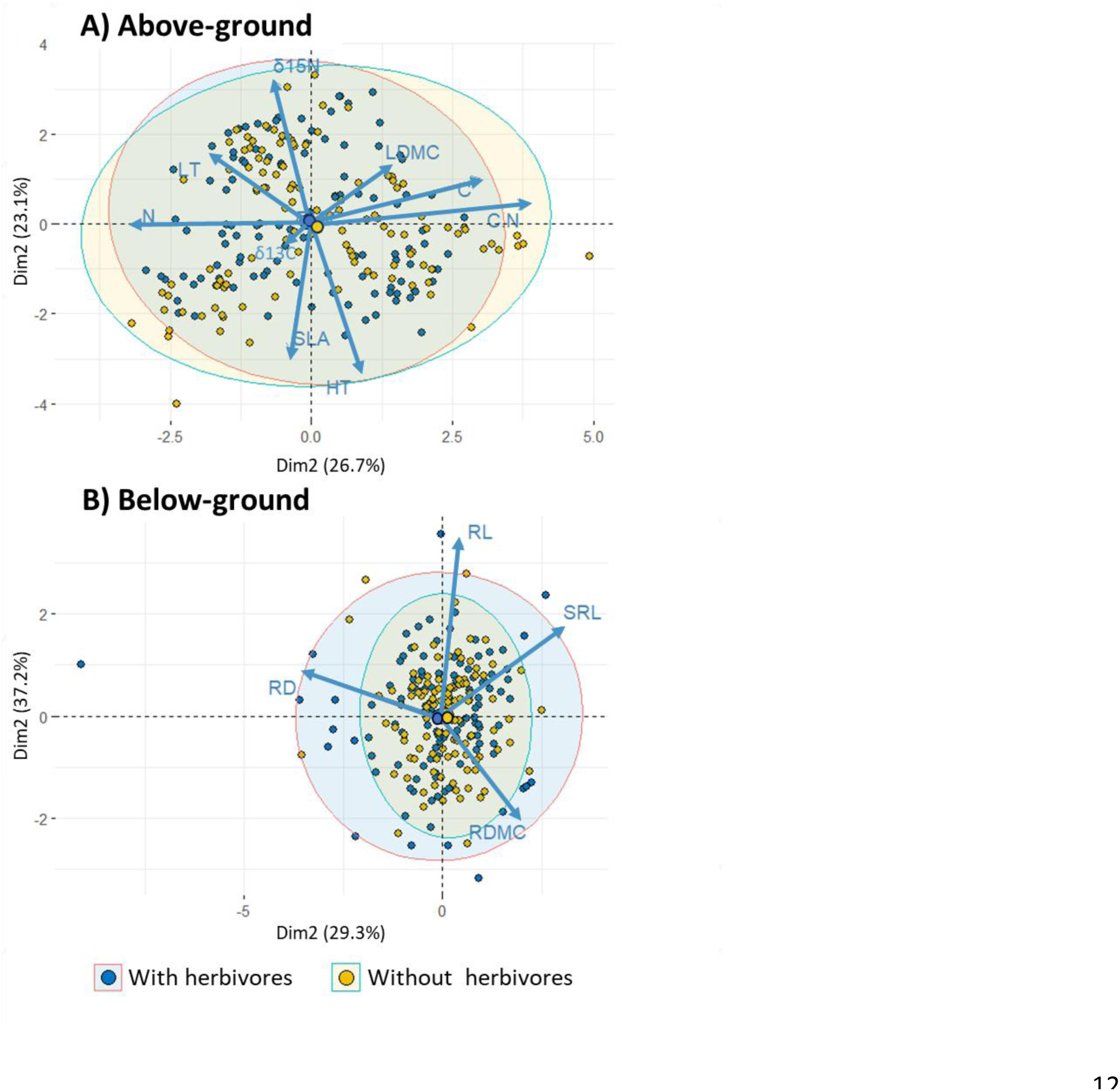
Trait spaces of: A) Above-ground and B) Below-ground plant organs of annual plant species in experimental treatments with and without herbivores in Bosque Fray Jorge National Park, Chile. The functional traits considered are height (HT), leaf thickness (LT), leaf dry matter content (LDMC), specific leaf area (SLA), leaf isotope 15 N (δ^15^N), leaf isotope 13 C (δ^13^C), leaf nitrogen content (%N), leaf carbon content (%C), leaf nitrogen to carbon ratio (CN), root dry matter content (RDMC), root diameter (RD), root length (RL) and specific root length (SRL). Projections are of the first two dimensions from the principal components analysis (PCA). Each symbol represents individual plants, with larger circles representing the centroids of each experimental treatment. The ellipses denote the 95% confidence intervals for treatments with and without herbivores.

The first two components of the PCA of below-ground traits explained 66.5.1% of the total variance (Fig. 2B; Tables S5 and S6). The first principal component explained 37.2 % of the variance and is strongly influenced by RD, which captures the collaboration gradient (Fig. 2B; Tables S5 and S6). Lower RD values (right) are consistent with the "do it yourself" strategy, while higher values (left) could indicate a stronger association with soil mycorrhizal fungi. The second principal component explained 29.3% of the variance and captured the below-ground resource acquisition gradient. This gradient is mainly associated with variation in RL (Fig. 2B, Table S6). Thus, longer roots indicate greater resource acquisition, and shorter roots indicate greater resource conservation (Fig. 2, Table S6). The PERMANOVA indicates statistically significant differences in trait spaces between species (P= 0.001) and treatments (P=0.004), but not for the interaction of both (P=0.159).

We observed differences in trait coordination between experimental treatments. In the treatment with herbivores, the above- and below-ground economic strategies (i.e., LES and RES) were significant and positively correlated (r=0.39, P=0.0001, Fig. 3, Table S6), yet in the treatment without herbivores they were not (r=0.05, P=0.58). When examining the coordination of traits for each species within each treatment, only *V. pusilla* exhibited coordination between above- and below-ground economic strategies, with a positive correlation in the herbivore treatment (Fig. 3, Table S6).

**Figure 3.**
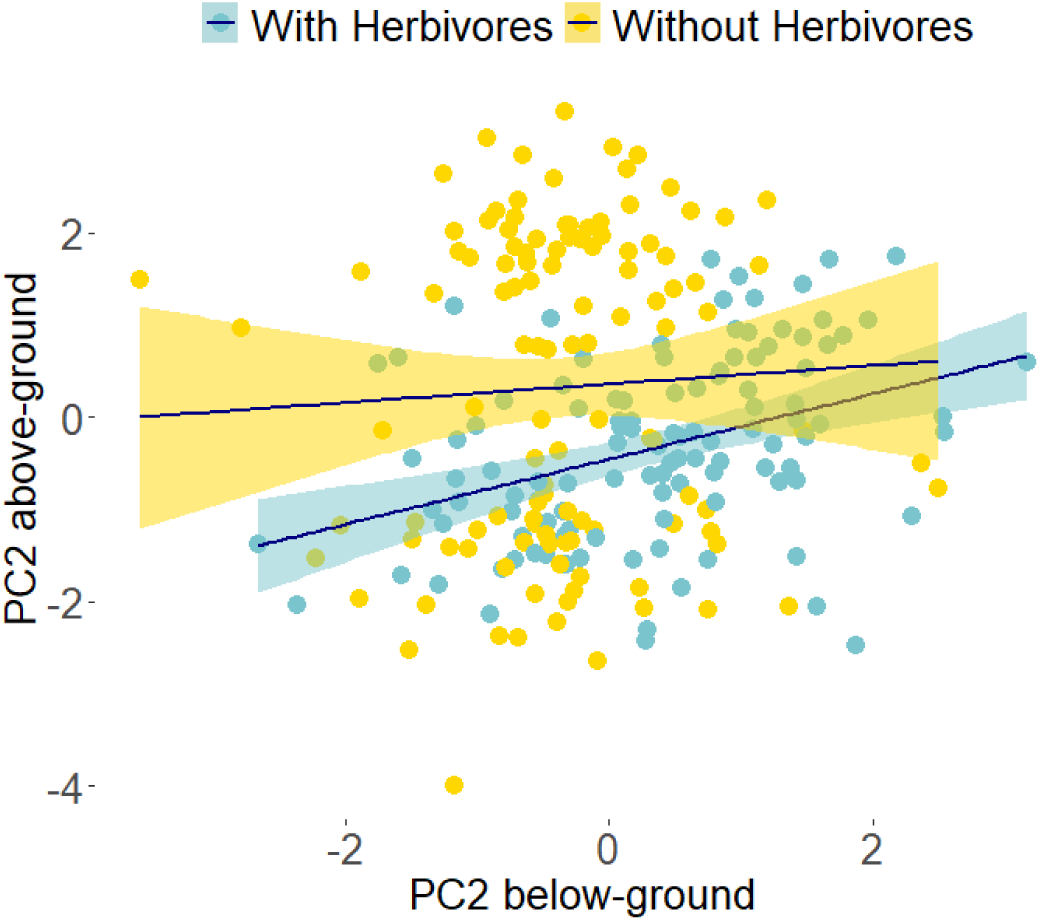
Correlation of above-ground and below-ground resource economics strategies. The Leaf Economics Spectrum (PC2 above-ground; LES) captures variation in HT, δ^15^N and SLA, while the Root Economics Spectrum (PC2 below-ground; RES) reflects variation in RL and RDMC. We compare the degree of co-ordination of above- and below-ground strategies between treatments, each of the points correspond to individuals of each species in each treatment.

### Response of above-ground ecological strategies to herbivores

We found shifts in dominant ecological strategies within and across species in response to the presence of herbivores. At the community level, plants growing with herbivores showed a higher mean tolerance to herbivores and higher variation and less kurtosis in herbivore tolerance than those in the treatment without herbivores (Fig. 4A, Table 1). In terms of intraspecific trait variation, individuals of *B. berteroanus* and *P. hispidula* growing with herbivores exhibited a higher mean tolerance to herbivores, as well as more skewness and more kurtosis than those growing without herbivores (Fig. 4C y 4G, Tables 1 and S8). Individuals of *M. pinnatifida* growing with herbivores also showed higher mean tolerance to herbivores and more variance in tolerance to herbivores, but less kurtosis and skewness than those growing without herbivores (Fig. 4E, Tables 1 and S8). Lastly, *V. pusilla* exhibited differences in tolerance to herbivores between treatments (Fig. 4I, Tables 1 and S8), with a higher mean and variance of tolerance to herbivores and lower kurtosis in the treatment with herbivores than in the treatment without herbivores.

**Figure 4.**
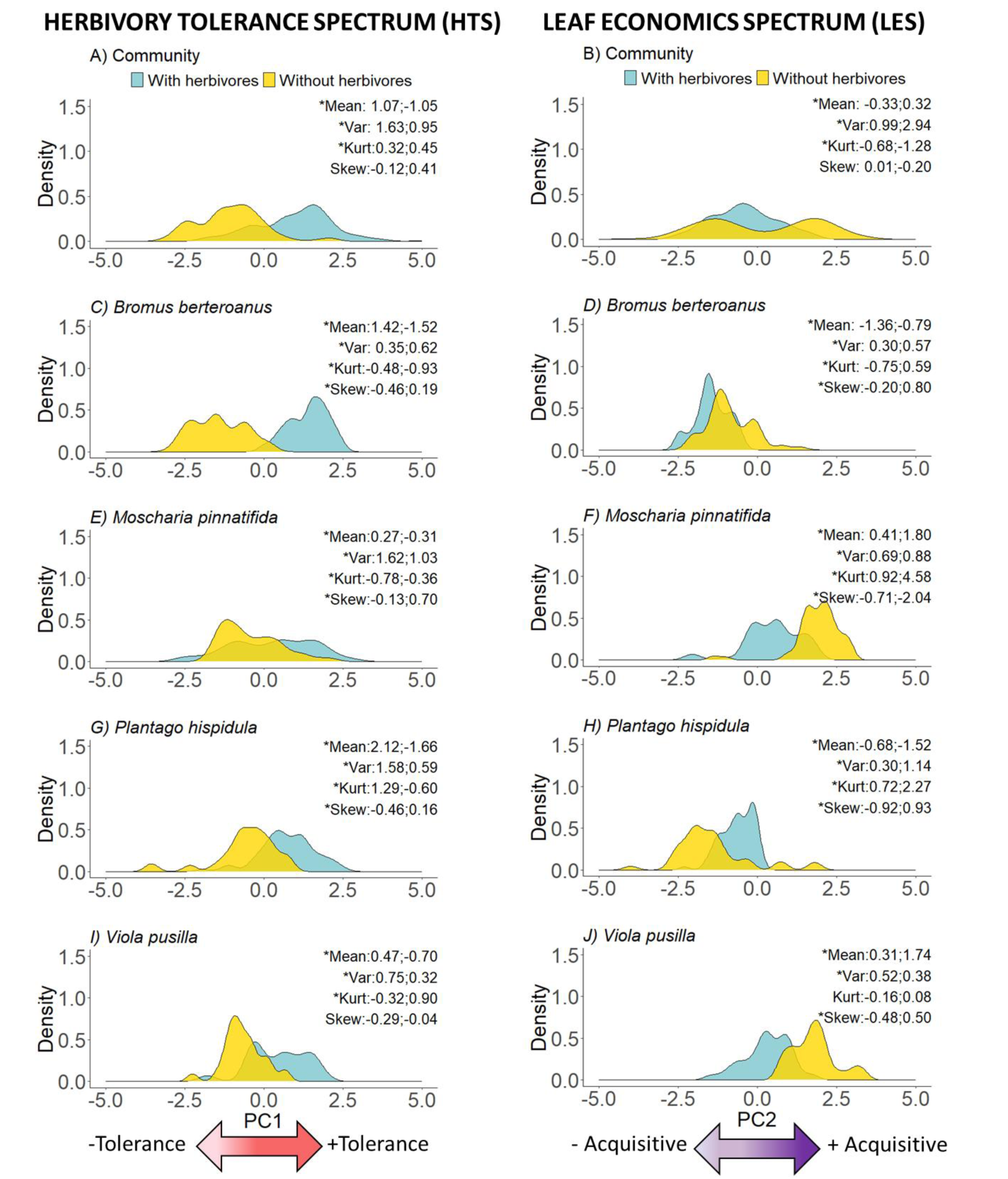
Variation in in two above-ground ecological strategies, HTS (left column corresponds to the PC1 of above-ground traits) and the LES (right column corresponds to the PC2 of above-ground traits) within and across annual species in a long-term herbivore exclusion experiment in Fray Jorge National Park, Chile at the community level (A & B) and at the species level (C – J). In figure insets, we provide mean values for each experimental treatments (with herbivores (blue); without herbivores (yellow)) for the following statistical moments of distributions of ecological strategies: mean, variance (var), kurtosis (Kurt) and skewness (Skew). We interpret differences between treatments as statistically significant if their 95% confidence intervals do not overlap with zero, which are indicated with an asterisk.

Annual plant communities growing with herbivores exhibited a more conservative resource acquisition strategy than those growing without herbivores (Fig. 4B, Table S8). Additionally, the resource acquisition strategy of annual plant communities growing in the presence of herbivores exhibited lower variation and higher kurtosis than that of annual plant communities growing without herbivores. Three of the four species exhibited shifts towards a more acquisitive resource-use strategy in the treatment without herbivores (Fig. 4D, H & J, Table S8). Individuals of *B. berteroanus* showed differences in the mean resource acquisition strategy between treatments, tending to be more acquisitive and showing greater variance, higher kurtosis and skewness in the treatment without herbivores than those in the treatment with herbivores (Fig. 4D, Table S8). For both *M. pinnatifida* and *V. pusilla*, individuals were decidedly more acquisitive in the treatment without herbivores than those in the treatment with herbivores (Fig. 4F & J, Table S8). *M. pinnatifida* exhibited greater variance, higher kurtosis and lower skewness of resource acquisition in the treatment without herbivores than in the treatment with herbivores (Fig. 4F, Table S8), while *V. pusilla* exhibited lower variance and a more skewed distribution of resource acquisition in the treatment without herbivores than in the treatment with herbivores (Fig. 4J, Table S8). In contrast to the other species, individuals of *P. hispidula* exhibited a more resource acquisitive strategy in the treatment with herbivores, yet lower variance, kurtosis and skewness than in the treatment without herbivores (Fig. 4H, Table S8).

### Response of below-ground ecological strategies to herbivores

Trait variation in experimental communities without herbivores was on average dominated by species with a more collaborative below-ground strategy than those in the treatment with herbivores (Fig. 5A, Table S6). We also observed differences in variance and kurtosis between treatments, with distributions of the CS in experimental communities without herbivores having higher variance and kurtosis than the experimental communities with herbivores. In terms of intraspecific variation, we observed differences in the mean, variance, skewness and kurtosis of the CS across treatments for *B. berteroanus*.

**Figure 5.**
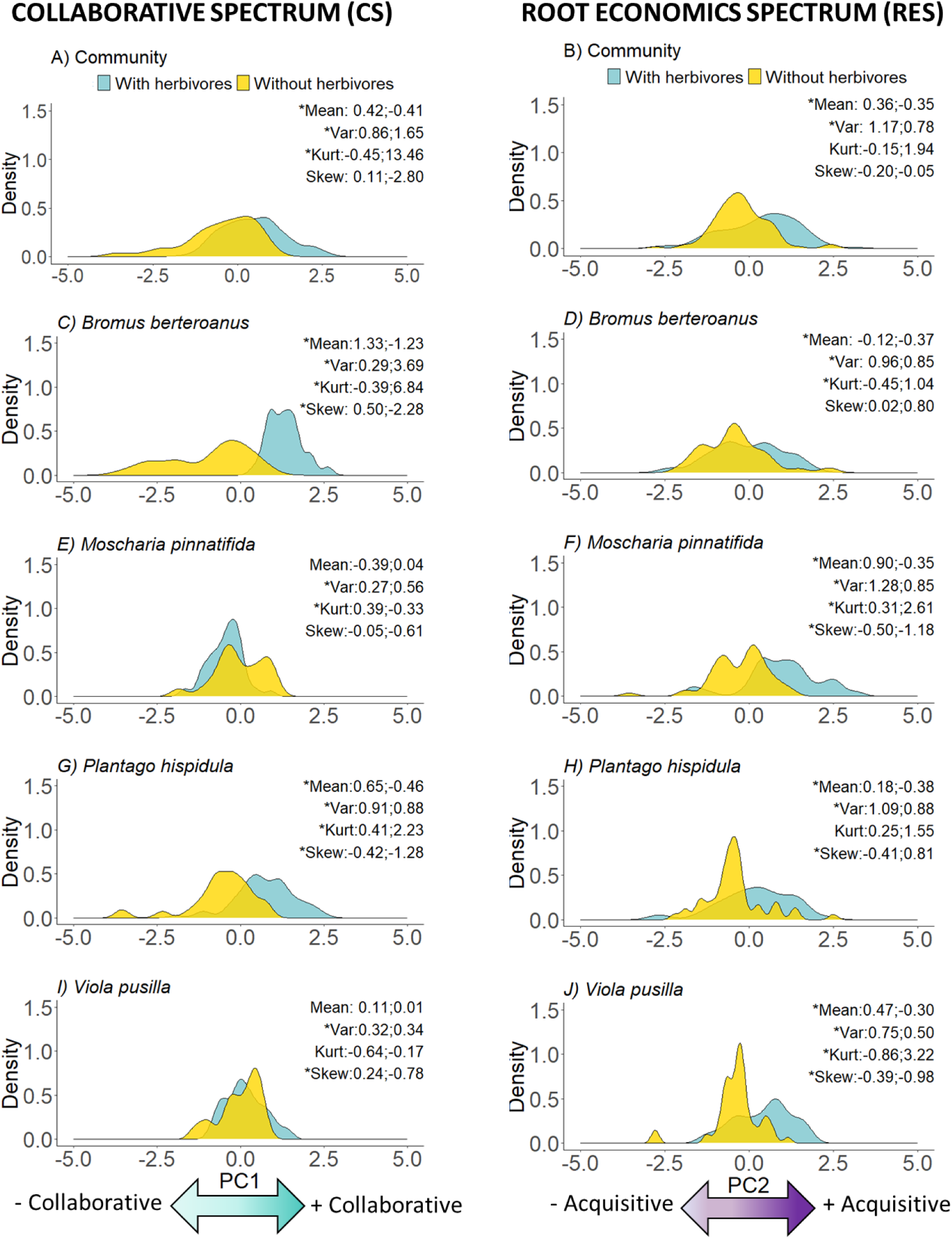
Variation in two below-ground ecological strategies, CS (left column corresponds to the PC1 of below-ground traits) and RES (right column corresponds to the PC2 of below-ground traits) within and across annual species in a long-term herbivore exclusion experiment in Fray Jorge National Park, Chile at the community level (A & B) and at the species level (C - J). In figure insets, we provide mean values for each experimental treatment (with herbivores (blue); without herbivores (yellow)) for the following statistical moments of distributions of ecological strategies: mean, variance(var), kurtosis (Kurt) and skewness (Skew). We interpret differences between treatments as statistically significant if their 95% confidence intervals do not overlap with zero, which are indicated with an asterisk.

Specifically, individuals in the treatment without herbivores were less collaborative on average, and had higher variation and kurtosis, and a more skewness than individuals in the treatment with herbivores (Fig. 5C, Table S7). Individuals of *M. pinnatifida* were more collaborative and had a lower variance and higher kurtosis in the treatment with herbivores than in the treatment without herbivores (Fig. 5E, Table S7). For *P. hispidula*, we found differences in mean values showed differences between treatments, indicating that individuals in the treatment without herbivores were more cooperative than those in the treatment with herbivores. We also found higher variance in this treatment, while both kurtosis and skewness were higher in the treatment without herbivores than in the treatment with herbivores (Fig. 5G, Table S7). Regarding *V. pusilla*, individuals in the treatment with herbivores had lower variance and higher skewness of the collaborative strategy than those in the treatment without herbivores (Fig. 5I, Table S7).

At the community level, we detected differences in the mean and variance among treatments for the RES. Species in the treatment without herbivores exhibited, on average, a more conservative resource acquisition strategy than those with herbivores. The distribution of the below-ground resource acquisition strategy exhibited a lower variance in the treatment without herbivores compared to the treatment with herbivores (Fig. 5B, Table S7). The comparison of the distribution of *B. berteroanus* reveals that individuals are slightly more conservative, with lower variation and higher kurtosis of resource acquisition in the absence of herbivores compared to those in the presence of herbivores (Fig. 5D, Table S7). *M. pinnatifida* and *V. pusilla* individuals in the treatment with herbivores had higher mean and variance and lower kurtosis and more symmetry of resource acquisition compared to those in the treatment without herbivores (Fig. 5F & J, Table S7). Finally, *P. hispidula* showed differences between treatments, with individuals growing with herbivores being less conservative, and having higher variance and lower symmetry compared to individuals in the treatment without herbivores (Fig 5 H. Table S7).

## DISCUSSION

We examined trait variation within and across annual plant species in response to herbivore loss. Our results reveal that a greater diversity of intraspecific ecological strategies may enhance species coexistence in the presence of herbivores (Figs. 4 and 5, Table S7). Across species, we found that the absence of herbivores leads to lower tolerance strategies, a more acquisitive above-ground resource acquisition strategy and a more conservative below-ground resource acquisition strategy, as well as more collaboration with soil microorganisms. In general, the herbivory tolerance spectrum (HTS) and below-ground resource economic spectrum (RES) exhibited lower variation in the absence of herbivores than the (above-ground) leaf economic spectrum (LES) and below-ground collaboration spectrum (CS), and LES, CS and RES had more skewed distributions in the absence of herbivores. Together, these changes in the diversity of ecological strategies within and across species may underpin how ecosystems will respond to biodiversity loss. Our results also highlight how herbivores appear to mediate coordination between above- and below-ground resource acquisition strategies, which weakens in the absence of herbivores (Fig 3, Table 6S).

### Trait responses to herbivore loss

Our study compared above- and below-ground trait responses in the presence and absence of herbivores within and among species. We find that community-level responses may differ from species-level responses. For instance, while three of the four studied species exhibited a more acquisitive strategy above-ground when growing without herbivores, as observed at the community level, *P. hispidula* exhibited a more conservative resource use strategy. These results demonstrate that different ecological strategies among species can provide a competitive advantage by differentiating the most abundant species, which could promote coexistence. (Chesson 2000). Moreover, the contrasting species-level ecological strategies observed in this study may contribute to enhancing the temporal stability of multiple ecosystem properties via compensatory dynamics (de Bello et al. 2021).

Our results further demonstrate that intraspecific variation may contribute to the successful establishment of dominant annual plants in the absence of herbivores. Our analysis of multiple statistical moments of distributions of ecological strategies allowed us to detect changes in the intra-specific trait diversity of dominant plant species in response to the absence of herbivores. For instance, the greater variance and lower skewness of *P. hispidula* than other species across most ecological strategies suggests that this species may have a high capacity to adapt to changes in competitive hierarchies associated with the loss of herbivores (Soliveres et al., 2015, Gallien 2017). Our results concur with those of other studies, which have shown the key role played by intraspecific variation in determining plant persistence and community assembly in semi-arid Mediterranean shrublands (Gross et al., 2013; Le Bagousse-Pinguet et al., 2015).

### Above-ground ecological strategies

Our study provides empirical evidence that the experimental exclusion of herbivores increases the diversity of above-ground resource acquisition strategies of annual plants within and across species (Fig. 4, right column). Three of the four species studied showed higher variation without herbivores, particularly *P. hispidula*, the most abundant annual species in our study (Fig. S2; Meserve et al., 2016; Fernandez-Murillo et al., 2023).

Additionally, we observed differences in the mean, variance, kurtosis, and skewness of the above-ground resource acquisition strategy (LES). The higher diversity of this ecological strategy observed in the treatment without herbivores may confer greater adaptability to diverse environmental conditions, thus providing a competitive advantage over species with lower LES diversity (Bennet et al., 2016; Mitchell et al., 2017). These findings support the idea that dominant plant species tend to exhibit a wider distribution of ecological strategies associated with resource acquisition, enabling them to adapt more readily to their surroundings (Adler et al., 2013).

Across species we find variation in traits associated with above-ground resource acquisition, particularly in the treatment without herbivores (Fig. 4). This variation can be primarily attributed to the diverse above-ground resource acquisition of *P. hispidula* and, to a lesser extent*, B. berteroanus* or *M. pinnatifida* (Fig. 4D, F, H). In a previous study within the same experiment, *B. berteroanus* and *P. hispidula* demonstrated the highest plant cover in the absence of herbivores (Fernandez-Murillo et al., 2023, see Fig. S2). This underscores the importance of these species in the absence of herbivores, as they exhibit trait values closely associated with resource acquisition, while the other two species (*M. pinnatifida* and *V. pusilla*) adopt a more conservative resource acquisition strategy. These findings suggest that the substantial variability in above-ground resource acquisition in the absence of herbivores could significantly influence competitive niche dynamics among dominant species (Maitner et al., 2023). At the intra-specific level, our findings show higher levels of skewness of the herbivore tolerance strategy in the treatment with herbivores, especially for *P. hispidula* and *B. berteroanus* (Fig. 4 C and G). These two species are essential components of the small mammal diet at our study site (Meserve et al., 1981; Kelt et al., 2013), and have maintained their dominance throughout the study period despite recent declines in precipitation (Guiterrez et al., 2010; Meserve et al., 2016; Fernández-Murillo et al., 2023). Furthermore, this result is broadly consistent with the expectations of “Trait Driver Theory” (Enquist et al., 2015), which postulates that skewness can be linked to community assembly processes such as competitive exclusion, suggesting that herbivores’ preference for species with lower leaf C has resulted in communities dominated by species with high leaf C and, therefore, greater herbivore tolerance.

### Below-ground ecological strategies

In contrast to expectations (e.g., Roumet et al. 2006; Foxx & Fort 2019), we observed a shift towards a more conservative below-ground resource acquisition strategy in response to the exclusion of herbivores. In scenarios of increased competition between plants, such as we assume to occur in the treatment without herbivores, we observed a change in the mean of the below-ground resource acquisition strategy, from a more acquisitive to a more conservative resource use strategy, resulting in shorter root lengths. As noted by Ravenek et al. (2016), not all root traits may confer a competitive advantage, and the duration of competition may be a critical factor in determining the relative importance of traits. Given our focus on annual plants, which are characterized by short life cycles and strong associations with mycorrhizal fungi (Roumet et al., 2006), we suggest that root length may be less important than root diameter (associated with the CS in this study; Fig. 5) in determining competition for below-ground resources. Thus, our results suggest that, in response to the increased competition among plants in the treatment without herbivores, annual plants may acquire below-ground resources more rapidly by investing resources in traits such as root diameter that strengthen associations with mycorrhizal fungi. While previous studies have shown the impacts of herbivore loss on above-ground competition (e.g., Fernández-Murillo et al., 2023), ours reveals that herbivore loss may trigger changes in how plants compete both above- and below-ground for critical resources in a semi-arid shrubland.

Contrary to our initial hypothesis, we observed that, across species, individuals depend on a collaborative strategy to acquire below-ground resources to a greater extent in the treatment without herbivores than in the treatment with herbivores (Bergmann et al., 2020; Weigelt et al., 2021). Additionally, at the intraspecific level, both *B. berteroanus* and *P. hispidula* displayed a tendency toward increased collaboration with mycorrhizal fungi in the treatment without herbivores, while *M. pinnatifida* and *V. pusilla* showed a trend towards values associated with lower levels of collaboration, although not statistically confirmed (Fig. 5). As cover of annual plants in the treatment without herbivores is significantly higher than in those with herbivores, where plant-plant competition is more intense (Meserve et al., 2016; Fernandez-Murillo et al., 2023), our results suggest that ecological strategies that contribute to efficient and rapid acquisition of below-ground resources - *via* plants’ association with mycorrhizal fungi - likely play a crucial role in determining competitive outcomes in our study system. Our findings are consistent with previous studies showing that plant species with mycorrhizal associations secure a larger portion of resources, giving them a competitive advantage in resource-limited environments such as semi-arid ecosystems (Delavaux et al., 2017; Ma et al., 2020; Han et al., 2022).

Our findings highlight a significant decrease in root diameter in the presence of herbivores, suggesting a reduced collaboration with soil mycorrhizal fungi (Fig. 5A). This pattern may be attributed to two main factors: Firstly, the presence of herbivores and associated fecal and urinary inputs has been shown to enhance the availability of soil organic matter and nutrients (Lemoine & Smich, 2019). Under such conditions, the symbiotic relationship between fungi and host plants is often weakened, as a plant’s contribution of nutrients to soil fungi becomes less essential because soil fungi may obtain nutrients from the surrounding environment (Frew et al., 2024). Secondly, the weaker collaboration with mycorrhizal fungi could be due to physical disruption caused by herbivores digging burrows, which may disrupt hyphal networks (Eldridge & Delgado-Baquerizo, 2018).

Previous studies have suggested that grazing by livestock and other large herbivores can decrease fungal hyphal length by removing plant material (Miller et al., 1995; Van der Heyde et al., 2017). The reduction in the plant-fungus relationship would imply less microbial activity, which could affect the productivity of the community (Li et al., 2022).

### Coordination of above- and below-ground resource economics strategies

Our findings suggest a potential shift in the coordination between above- and below- ground plant traits linked to resource acquisition in response to herbivore exclusion (Fig. 3). In the presence of herbivores, a potential coordination between above- and below- ground ecological strategies is observed, which is lost or relaxed in the absence of herbivores. We attribute the decoupling of above- and below-ground resource acquisition strategies with the influence of small mammals on plant consumption (Gutierrez & Meserve 2000) and their indirect effects on soil structure through burrow construction (Laundré & Reynolds 2013; Louw et al., 2019) and surface disturbance as they search for food (Brown & Heske 1990) and soil nutrients (Poe et al., 2019). These changes in abiotic conditions trigger a shift in the dominant processes shaping community assembly of annual plants, from herbivory (treatment with herbivores) to competition (treatment without herbivores). While the exclusion of herbivores, by allowing an increase in plant biomass, may stabilize soils and reduce soil water loss, it also intensifies competition for other resources, such as space, light, and nutrients (Weiner & Thomas et al., 1992; Naidu et al., 2021; Eskelinen et al., 2022; Roy & Bagchi 2022). Our results suggest that the decoupling of above- and below-ground resource acquisition strategies may arise from changes in underlying ecological processes associated with the loss of herbivores.

Our results suggest that trait coordination at the local scale may respond primarily to biotic factors rather than abiotic factors such as temperature or precipitation. Previous studies in Mediterranean ecosystems have shown that resource acquisition or conservation is a coordinated response observed throughout the plant (e.g., Liu et al., 2010). Our results from control conditions (e.g., with herbivores) are consistent with these observations, where such coordination is evident (see Fig. 3B). In contrast, other studies have suggested that the coordination of above- and below-ground resource acquisition strategies is the result of increased intra-specific variation attributed to crowding or neighborhood effects (Le Bagousse-Pinguet et al., 2015, Afesa et al., 2022). This appears consistent with the coordination observed between above- and below-ground resource acquisition strategies in our herbivore exclusion treatment. Under such conditions, higher plant biomass may lead to crowding and to greater intra-specific variation above- than below-ground (see Fig. 5, right column).

Within species, the coordination of resource acquisition strategies both above- and below- ground across treatments appeared to be more subtle. Among the four species studied here, only *V. pusilla* exhibited a marked coordination of above- and below-ground traits (Table S6). For *V. pusilla* in the herbivore treatment, resource acquisition strategies appeared to be closely aligned as anticipated, indicating that individuals displaying acquisitive behavior above-ground tended to exhibit similar traits below-ground. Such coordination is commonly associated with a potential competitive advantage (Reich et al.,2014, Foxx & Ford et al., 2019). However, the observed divergence in above- and below-ground strategies among the other three species (Table S6) may not necessarily signify a disadvantage. Instead, it could suggest the potential for a more efficient resource allocation by individuals in environments with limited resources (Bergmann et al., 2016). This also suggests that, while herbivores might not directly alter a species’ resource acquisition strategy, individuals not experiencing herbivore pressure might reallocate resources towards competition with conspecific and heterospecific individuals (Zhou et al., 2018), as evidenced by observed shifts in the collaboration gradient and herbivory tolerance at the intraspecific level.

## CONCLUSIONS

The long-term exclusion of vertebrate herbivores at our field site in north-central Chile appears to have induced shifts within and across species in the diversity of ecological strategies employed by annual plants. In particular, plant species grown in the absence of vertebrate herbivory are associated with weaker collaborations with soil microorganisms and lower herbivore tolerance, and exhibit more acquisitive resource acquisition strategies, relative to plants exposed to herbivory. These results suggest that the loss of herbivores in this ecosystem may fundamentally alter how plants compete for limited above- and below-ground resources. By changing competitive niche dynamics, these observations suggest that the loss of such herbivores has the potential to restructure annual plant communities.

## Supporting information

Information_Support

## ACKNOWLEDGMENTS

This work was possible thanks to the vegetation team (2015-2021) of the long-term research project LTSER Fray Jorge, particularly Víctor Pastén and Gerardo Gutiérrez. The grants that supported this work were FONDECYT 1220358 (FDA), 1201347 (DC), 10302225, and 1160026, as well as NSF LTREB-DEB2025816 (DAK), ACE210006, and PIA/BASAL FB210006.

## AUTHOR CONTRIBUTIONS

MPFM, FA, and DC conceived the idea and developed the objectives and hypotheses. MPFM and FA developed the methodology. MPFM and DC analyzed the data. MPFM and AT collected plant samples in the field. JRG, DK, and PM founded and oversee the LTSER Fray Jorge. All authors contributed to the writing, drafting, and revision of the article. Our research unites authors from various countries, encompassing scientists situated in the nation where the study was conducted.

## DATA AVAILABILITY STATEMENT

Data available from: https://figshare.com/s/6febdf5f1c2b8c4e2951

## SUPPORTING INFORMATION

**Table S1.** List of traits measured in annual plants.

**Table S2.** Variance explained by two principal components (PC) for above-ground functional traits.

**Table S3.** Loadings of the functional traits onto the first two principal components obtained from PCA of above-ground functional traits.

**Table S4.** Variance explained by two principal components (PC) for below-ground functional traits.

**Table S5**. Loadings of variables onto the first three principal components obtained from the PCA of below-ground traits.

**Table S6.** Pairwise correlations of across and within-species above- and below-ground resource acquisition strategies.

**Table S7.** Results of the stratified bootstrapping analysis conducted on the first two dimensions of the PCA focusing on above- and below-ground traits.

**Figure S1**. Mean annual precipitation in LTSER Bosque Fray Jorge NP.

**Figure S2.** Coverage of dominant annual plants species in 2021.

