## Supplementary material for "Within and Across Species Variation in Plant Traits in Response to Long-Term Experimental Herbivore Exclusion in a Semi-Arid Shrubland": Information_Support

Authors: María del Pilar Fernández-Murillo[1,2,3]; Fernando Daniel Alfaro[2]; Douglas A. Kelt[5]; Peter L. Meserve[6]; Julio R. Gutiérrez[3†], Alejandra J. Troncoso[3,4] & Dylan Craven[2,7]
Article acceptance date: Click here to enter a date.

The following Supporting Information is available for this article:

**Table S1. List of traits measured in annual plants**

List of traits measured in annual plants of LTERB National Park Bosque Fray Jorge with their abbreviations, units, definition, ecological strategy and references.

| Trait | Abbr | Units | Definition | Ecological strategy | Reference |
|---|---|---|---|---|---|
| Specific Leaf Area | SLA | $cm^2 g^{-1}$ | Leaf area/leaf dry mass | +Acquisitive | Hodgson et al., 2011, Zheng et al., 2015 |
| Height | HT | mm | Maximum vegetative plant height | +Acquisitive | Diaz et al., 2001; Cingaloni et al.,2005 |
| Leaf dry matter content | LDMC | $gg^{-1}$ | Leaf dry mass/fresh mass | - Acquisitive | Hodgson et al., 2011 |
| Leaf thickness | LT | mm | Leaf thickness | -Acquisitive | Hodgson et al., 2011 |
| Specific root length | SRL | $mg^{-1}$ | Root length/dry mass | -Collaboration | Bergmann et al., 2000; Weigelt et al., 2021 |
| Root dry matter content | RDMC | $gg^{-1}$ | Root dry mass/fresh mass | - Acquisitive | Kramer-Walter et al., 2016 |
| Root diameter | RD | mm | Average root diameter | +Collaboration | Bergmann et al. 2000; Weigelt et al., 2021 |
| Total Root Length | RL | mm | Average root length | + Acquisitive | Kramer-Walter et al., 2016 |
| Delta 15 N | $\delta^{15}N$ | | $^{15}N$ isotopic abundance in leaves | - Tolerance | Diaz et al., 2007 |
| Delta 13 C | $\delta^{13}C$ | | $^{13}C$ isotopic abundance in leaves | + Tolerance | Diaz et al., 2007 |

| | | | | | |
|---|---|---|---|---|---|
| N | %N | % | Leaf nitrogen concentration | - Tolerance | Diaz et al., 2007; Carmona et al., 2011; Zvereva & Kozlov 2014; Weigelt et al., 2021 |
| C | %C | % | Leaf carbon concentration | + Tolerance | Diaz et al., 2007; Carmona et al., 2011; Zvereva & Kozlov 2014; Weigelt et al., 2021 |
| C/N | C/N | | Leaf ratio carbon to nitrogen | +Tolerance | Carmona et al. 2011 |

**Table S2. Variance explained by two principal components (PC) for above-ground functional traits**

Variance explained by two principal components (PC) for above-ground functional traits of annual plants in the Fray Jorge LTSER. Variance and Percentage of Variance (% of var.) are provided for each principal component. Additionally, the Cumulative Percentage of Variance (Cumulative % of var.) is shown for each principal component."

| | **PC.1** | **PC.2** |
|---|---|---|
| Variance | 2.41 | 2.08 |
| % of var. | 26.73 | 23.12 |
| Cumulative% of var | 26.73 | 49.85 |

**Table S3. Loadings of the functional traits onto the first two principal components obtained from PCA of above-ground functional traits.**

Loadings of the functional traits onto the first two principal components (PC) obtained from the principal component analysis (PCA) of above-ground functional traits of annual plants in the Fray Jorge LTER. HT refers to plant height, LT refers to leaf thickness, LDMC refers to leaf dry matter content, SLA refers to specific leaf area, $\delta^{15}N$ refers to the nitrogen isotope ratio, $\delta^{13}C$ refers to the carbon isotope ratio, N refers to nitrogen content, C refers to carbon content, and C/N refers to the carbon-to-nitrogen ratio. 'Ctr' represents the contribution of each trait to the respective principal component.

| | PC.1 | ctr | PC.2 | Ctr |
|---|---|---|---|---|
| HT | 0.21 | 1.90 | -0.81 | 31.58 |
| LT | -0.44 | 8.08 | 0.37 | 6.68 |
| LDMC | 0.34 | 4.87 | 0.31 | 4.71 |

| | | | | |
|---|---|---|---|---|
| SLA | -0.09 | 0.38 | -0.70 | 23.61 |
| $\delta^{15}N$ | -0.16 | 1.06 | 0.79 | 29.81 |
| $\delta^{13}C$ | -0.05 | 0.14 | -0.04 | 0.10 |
| N | -0.77 | 24.84 | -0.01 | 0.002 |
| C | 0.73 | 22.42 | 0.25 | 2.97 |
| C/N | 0.93 | 36.30 | 0.11 | 0.55 |

**Table S4. Variance explained by two principal components (PC) for below-ground functional traits.**

Variance explained by two principal components (PC) for below-ground functional traits of annual plants in the Fray Jorge LTER. Variance and Percentage of Variance (% of var.) are provided for each principal component. Additionally, the Cumulative Percentage of Variance (Cumulative % of var.) is shown for each principal component."

| | PC.1 | PC.2 |
|---|---|---|
| Variance | 1.48 | 1.17 |
| % of var. | 37.22 | 29.29 |
| Cumulative of var. % | 37.22 | 66.51 |

**Table S5. Loadings of variables onto the first three principal components obtained from the PCA of below-ground traits.**

Loadings of variables onto the first three principal components (PC) obtained from the principal component analysis (PCA) of below-ground traits of annual plants in the Fray Jorge LTER. 'Ctr' represents the contribution of each variable to its respective principal component.

| | PC.1 | Ctr | PC.2 | Ctr |
|---|---|---|---|---|
| RDMC | 0.52 | 18.51 | -0.41 | 14.30 |
| RD | -0.88 | 51.92 | 0.09 | 0.80 |
| RL | -0.11 | 0.89 | 0.80 | 54.21 |
| SRL | 0.65 | 28.69 | 0.60 | 30.69 |

**Table S6. Pairwise correlations of across and within-species above- and below-ground resource acquisition strategies.**

Pairwise correlations of across and within-species above- and below-ground resource acquisition strategies. The treatments without herbivore and with herbivore are indicated, and the studied species are *Bromus berteroanus* (Brbe), *Moscharia pinnatifida* (Mopi), *Plantago hispidula* (Plhi), and *Viola pusilla* (Vipu). The values in the Spearman correlation coefficient column represent the type of relationship (positive or negative) between above- and below-ground acquisition strategies, while the p-values indicate the significance of these relationships. Values <0.05 are highlighted in bold to denote significant correlations.

| | Spearman | P |
|---|---|---|
| **with herbivores** | **0.39** | **0.0001*** |
| without herbivores | 0.05 | 0.58 |
| Brbe with herbivores | 0.15 | 0.44 |
| Brbe without herbivores | -0.03 | 0.84 |
| Mopi with herbivores | -0.006 | 0.97 |
| Mopi without herbivores | 0.06 | 0.75 |
| Plhi with herbivores | 0.27 | 0.16 |
| Plhi without herbivores | 0.20 | 0.29 |
| **Vipu with herbivores** | **0.60** | **0.005*** |
| Vipu without herbivores | -0.16 | 0.40 |

**Table S7. Results of the stratified bootstrapping analysis conducted on the first two dimensions of the PCA focusing on above- and below-ground traits.**

Results of the stratified bootstrapping analysis conducted on the first two dimensions of the Principal Component Analysis (PCA) focusing on above- and below-ground traits. The values provided indicate confidence intervals derived from the percentiles obtained through the analysis with herbivores and without herbivores treatments. The analyses are conducted across various ecological levels, encompassing interspecific (community) and intraspecific levels. The investigation encompasses four species: *B. berteroanus*, *M. pinnatifida*, *P. hispidula*, and *V. pusilla*. The assessment encompasses four statistical moments: mean, variance (var), kurtosis (Kurt), and skewness (Skew). Confidence intervals that do not encompass zero are emphasized in bold, indicating the presence of statistically significant differences between treatments.

| | | Tolerance Herbivores | | | | LES | | | |
|---|---|---|---|---|---|---|---|---|---|
| | | with herbivores | | without herbivores | | with herbivores | | without herbivores | |
| | Moments | CI_low | CI_high | CI_low | CI_high | CI_low | CI_high | CI_low | CI_high |
| Community | mean | **0.98** | **1.16** | **-1.11** | **-0.98** | **-0.40** | **-0.26** | **0.20** | **0.44** |
| | var | **1.44** | **1.81** | **0.84** | **1.07** | **0.91** | **1.07** | **2.76** | **3.12** |
| | skew | -0.31 | 0.07 | 0.22 | 0.59 | -0.10 | 0.12 | -0.32 | -0.07 |
| | kurt | **0.02** | **0.63** | **0.06** | **0.83** | **-0.82** | **-0.54** | **-1.42** | **-1.13** |
| *B. berteroanus* | mean | **1.38** | **1.47** | **-1.57** | **-1.47** | **-1.40** | **-1.32** | **-0.85** | **-0.74** |
| | var | **0.32** | **0.38** | **0.58** | **0.67** | **0.27** | **0.32** | **0.51** | **0.64** |
| | skew | **-0.58** | **-0.34** | **0.09** | **0.30** | **-0.29** | **-0.11** | **0.65** | **0.94** |

| | | | | | | | | | |
|---|---|---|---|---|---|---|---|---|---|
| | kurt | **-0.69** | **-0.27** | **-1.05** | **-0.81** | **-0.88** | **-0.62** | **0.24** | **0.93** |
| *M. pinnatifida* | mean | **0.19** | **0.36** | **-0.39** | **-0.24** | **0.35** | **0.47** | **1.73** | **1.86** |
| | var | **1.49** | **1.75** | **0.94** | **1.12** | **0.61** | **0.78** | **0.73** | **1.04** |
| | skew | **-0.24** | **-0.02** | **0.59** | **0.81** | **-0.87** | **-0.56** | **-2.19** | **-1.88** |
| | kurt | **-0.90** | **-0.65** | **-0.59** | **-0.13** | **0.57** | **1.27** | **3.30** | **5.87** |
| *P. hispidula* | mean | **2.04** | **2.21** | **-1.72** | **-1.61** | **-0.72** | **-0.64** | **-1.60** | **-1.45** |
| | var | **1.37** | **1.79** | **0.54** | **0.64** | **0.26** | **0.33** | **0.98** | **1.31** |
| | skew | **-0.68** | **-0.23** | **0.06** | **0.26** | **-1.06** | **-0.77** | **0.71** | **1.15** |
| | kurt | **0.98** | **1.61** | **-0.75** | **-0.44** | **0.34** | **1.10** | **1.68** | **2.85** |
| *V. pusilla* | mean | **0.40** | **0.53** | **-0.74** | **-0.66** | **0.26** | **0.36** | **1.70** | **1.79** |
| | var | **0.68** | **0.82** | **0.28** | **0.36** | **0.47** | **0.57** | **0.34** | **0.42** |
| | skew | -0.43 | -0.14 | -0.26 | 0.17 | **-0.61** | **-0.35** | **0.40** | **0.61** |
| | kurt | **-0.52** | **-0.11** | **0.60** | **1.20** | -0.39 | 0.07 | -0.18 | 0.33 |
| | | Collaboration | | | | RES | | | |
| Community | mean | **0.36** | **0.49** | **-0.50** | **-0.32** | **0.28** | **0.44** | **-0.41** | **-0.29** |
| | var | **1.29** | **1.37** | **-1.36** | **-1.10** | **-0.19** | **-0.05** | **-0.44** | **-0.31** |
| | skew | -0.43 | -0.35 | -0.01 | 0.09 | 0.83 | 0.98 | -0.41 | -0.28 |
| | kurt | **0.58** | **0.71** | **-0.53** | **-0.40** | 0.10 | 0.25 | -0.45 | -0.31 |
| *B. berteroanus* | mean | **0.07** | **0.15** | **-0.04** | **0.04** | **0.41** | **0.53** | **-0.35** | **-0.25** |
| | var | **0.78** | **0.93** | **1.13** | **2.16** | **1.06** | **1.28** | **0.67** | **0.89** |
| | skew | **0.26** | **0.31** | **2.94** | **4.44** | 0.88 | 1.05 | 0.74 | 0.95 |
| | kurt | **0.24** | **0.30** | **0.51** | **0.61** | **1.14** | **1.42** | **0.73** | **0.98** |
| *M. pinnatifida* | mean | 0.81 | 1.01 | 0.76 | 1.01 | **0.97** | **1.22** | **0.76** | **1.00** |
| | var | **0.29** | **0.35** | **0.31** | **0.38** | **0.69** | **0.81** | **0.42** | **0.59** |
| | skew | -0.02 | 0.24 | -3.61 | -1.98 | **-0.36** | **-0.04** | **-0.38** | **0.28** |
| | kurt | **0.38** | **0.62** | **-2.46** | **-2.09** | **-0.10** | **0.15** | **0.65** | **0.95** |
| *P. hispidula* | mean | **-0.23** | **0.13** | **-0.73** | **-0.50** | **-0.61** | **-0.39** | **-1.39** | **-0.97** |
| | var | **-0.57** | **-0.28** | **-1.43** | **-1.13** | **-0.57** | **-0.25** | **0.63** | **0.99** |
| | skew | **0.14** | **0.35** | **-0.89** | **-0.67** | **-0.49** | **-0.29** | **-1.24** | **-0.72** |
| | kurt | **-0.63** | **-0.26** | **7.25** | **19.68** | -0.40 | 0.10 | 1.35 | 2.54 |
| *V. pusilla* | mean | -0.62 | -0.17 | 5.67 | 8.01 | **-0.60** | **-0.29** | **0.66** | **1.41** |
| | var | **0.16** | **0.62** | **-0.56** | **-0.09** | **0.02** | **0.60** | **2.07** | **3.15** |
| | skew | **0.14** | **0.68** | **1.65** | **2.80** | **-0.01** | **0.51** | **1.07** | **2.04** |
| | kurt | -0.80 | -0.48 | -0.45 | 0.10 | **-1.02** | **-0.70** | **2.64** | **3.80** |

**Figure S1**. **Mean annual precipitation in LTSER Bosque Fray Jorge NP.**

Mean annual precipitation in millimeters of the semiarid scrubland of the Bosque Fray Jorge National Park, the continuous line indicates the mean over the 33 years of measurement.

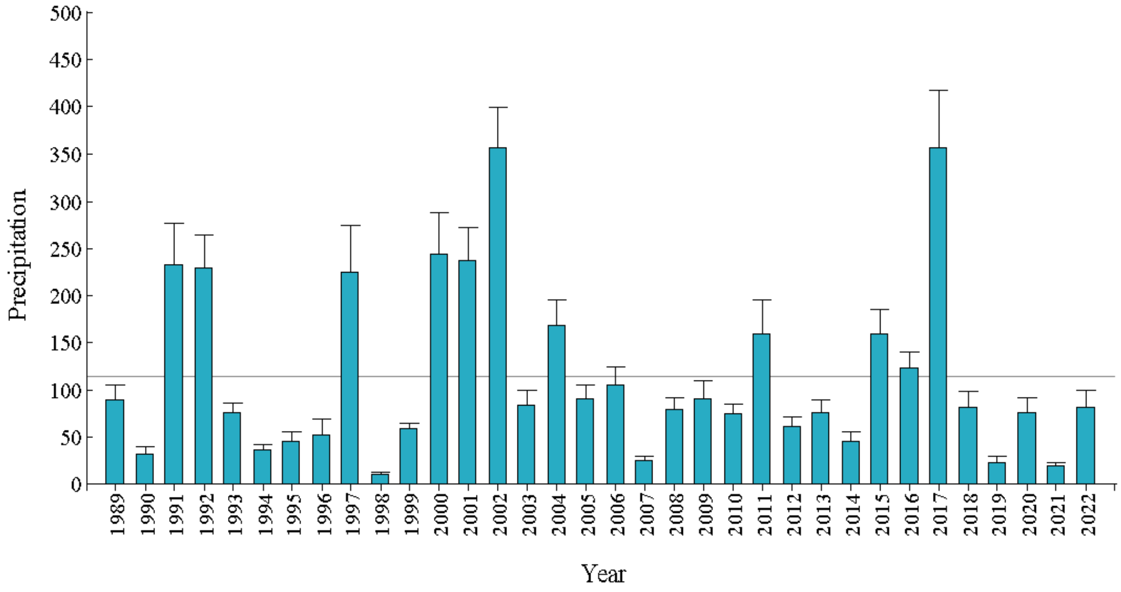


**Figure S2. Coverage of dominant annual plants species in 2021.**

(A-D) Dominant species. E) Coverage (%) of dominant species in 2021 in LTER Bosque Fray Jorge National Park. Unpublished data corresponding to the annual monitoring of the LTER Fray Jorge project, more information on the methodology in Meserve et al. 2016. Abbreviations correspond to dominant species: *Viola pusilla* (Vipu), *Bromus berteroanus* (Brbe), *Moscharia pinnatifida* (Mopi), *Plantago hispidula* (Plhi), and other species which may include geophyte species (Others) for the two treatments: with and without herbivores (blue and yellow respectively).

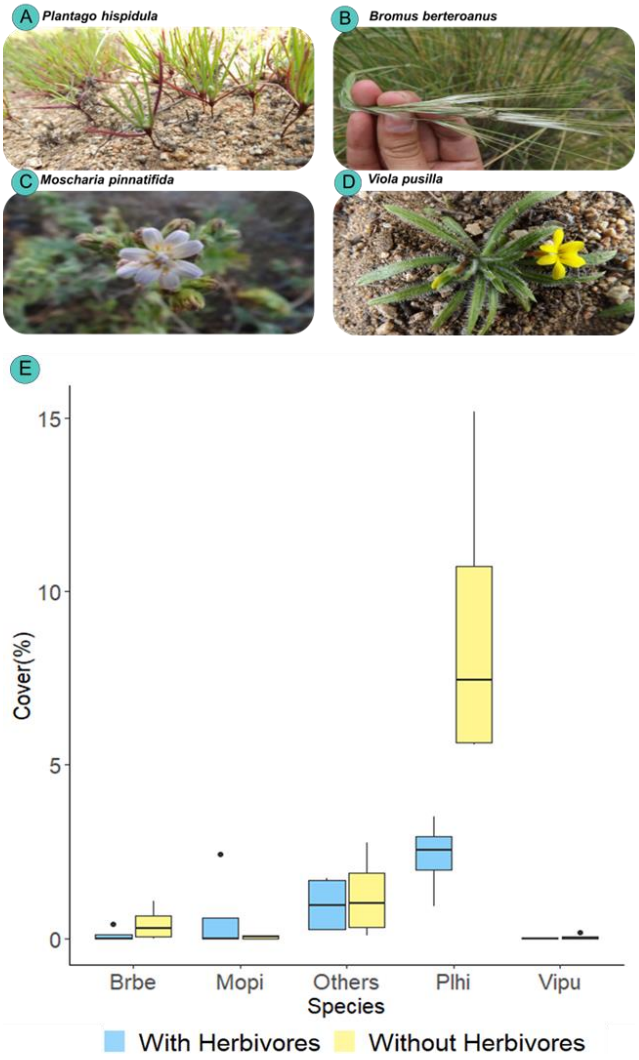

A Plantago hispidula
B Bromus berteroanus
C Moscharia pinnatifida
D Viola pusilla
E
Cover(%)
15
10
5
0
Brbe
Mopi
Others
Plhi
Vipu
Species
With Herbivores
Without Herbivores
